# Neurodevelopment Trajectory of Electrophysiological Functional Connectivity Following Alcohol Use Initiation

**DOI:** 10.64898/2026.06.20.733186

**Authors:** Alberto del Cerro-León, Danylyna Shpakivska-Bilan, Marcos Uceta, Fernando Maestú, Luis M García-Moreno, Luis Fernando Antón-Toro

## Abstract

**Background:** Adolescence is characterized by profound neurodevelopmental changes that shape large-scale brain network organization and may confer vulnerability to risk-taking behaviors, including alcohol use. While cross-sectional and prospective studies have examined functional connectivity (FC) alterations before and after consumption, there is little evidence of how networks evolve during adolescence.

**Methods:** The present longitudinal study investigated electrophysiological FC trajectories during alcohol initiation using resting-state magnetoencephalography (MEG). 61 alcohol-naïve adolescents (mean age at baseline = 14.4) were assessed and re-evaluated two years later (mean age = 16.4).

**Results:** At baseline, stronger FC in theta (4-8 Hz), alpha (8-12 Hz), and high-beta (20-30 Hz) bands predicted greater alcohol consumption at follow-up, replicating previous findings. Longitudinal analyses with linear mixed-effects models revealed significant stage × SAUs interactions across all three frequency bands. Adolescents with low-to-moderate alcohol use showed normative increases in FC over time, consistent with typical neurodevelopmental maturation. In contrast, heavier drinkers exhibited stabilization or reduction of FC, suggesting a divergence from normative trajectories. Notably, theta-band hyperconnectivity persisted after alcohol initiation and remained positively associated with current alcohol consumption, particularly across anteroposterior connections.

**Conclusion:** These findings indicate heterogeneous neurodevelopmental trajectories associated with alcohol use severity. Elevated pre-consumption connectivity, especially in the theta band, may reflect a vulnerability marker rather than solely a consequence of alcohol exposure. Overall, results highlight the importance of considering individual variability in brain maturation when examining adolescent substance use and suggest that early hyperconnectivity may signal increased risk for heavier alcohol involvement.

## 1. Introduction

Adolescence is a period of life during which the brain undergoes significant maturational changes that shape the development of essential cognitive functions and behavior to adapt to the psychosocial environment (1,2). Following the onset of puberty, a cascade of neurobiological processes promotes earlier structural and functional maturation of the mesolimbic dopaminergic pathway, which regulates motivational and reward-oriented behaviors, and a delayed development of the mesocortical circuit, associated with cognitive and behavioral executive control(2,3). This maturational imbalance between reward-seeking and executive mechanisms gives rise to distinctive behavioral traits during adolescence, characterized by heightened risk-taking tendencies, such as the initiation in alcohol consumption (4,5).

Alcohol consumption as a leisure activity is widely accepted in our society and has become progressively normalized among adolescents and young adults (6). According to the latest European School Survey Project on Alcohol and Other Drugs (ESPAD) data, nearly three-quarters of 15–16-year-olds in Europe report having consumed alcohol at least once, with around 40% reporting use in the past month. Approximately 30% engage in heavy episodic drinking, underscoring the high prevalence of alcohol use in mid-adolescence. These trends are particularly concerning given that the adolescent brain is particularly vulnerable to the neurotoxic effects of alcohol (7). Cross-sectional studies have consistently reported structural and functional alterations in young individuals who engage in heavy drinking (8,9). However, prospective studies has also revealed pre-existing structural (10,11) and electrophysiological (12,13) differences that precede alcohol initiation and are associated with the emergence of drinking behaviors years later. In previous studies conducted by our research group, prospective analyses showed that stronger coupling and activation across multiple brain networks and frequency bands during the resting state and cognitive tasks predicted heavier alcohol consumption two years later (12–15). Given this scenario, longitudinal studies are needed to enable us to separate predisposing traits from the effects of alcohol consumption on the brain. Nevertheless, evidence on how functional networks maturate longitudinally after drinking onset is scarce. To date, some studies have pointed out that normative development can be disrupted by the onset of heavy drinking, leading to reduced network segregation and abnormal frontal–occipital coupling (16). Structural MRI evidence further supports this disruption, showing that early alcohol use accelerates cortical thinning and attenuates white matter expansion, particularly in frontal regions (7). These alterations suggest that alcohol exposure may interfere with the neurodevelopmental mechanisms underlying functional integration and executive control. However, how electrophysiological functional networks (M/EEG), which index the temporal and frequency-specific dynamics of neurodevelopment, evolve longitudinally after alcohol use onset remains unexplored.

The present study aimed to characterize the evolution of electrophysiological functional connectivity trajectories following the onset of alcohol use. To this end, we assessed resting-state electrophysiological activity using MEG in a cohort of alcohol-naïve adolescents aged 13 to 15 years. After a two-year follow-up, participants were re-evaluated with the same MEG protocol, and their alcohol use habits were recorded. Changes in functional connectivity between both time points were analyzed as a function of alcohol consumption, controlling for biological sex and age. Previous work using pre-consumption data reported robust associations between higher functional connectivity and subsequent alcohol use. The current study extends these findings by disentangling the longitudinal evolution of these hyperconnectivity profiles. Based on prior literature on functional maturation and alcohol-related structural alterations, we hypothesized that adolescents with higher alcohol use would exhibit divergent trajectories of functional connectivity change between pre- and post-consumption stages.

## 2. Materials & Methods

### 2.1 Participants

The participants were recruited from high schools in the Madrid area as part of two longitudinal research projects funded by the Spanish Ministry of Health. Both projects followed the same evaluation protocol, which included two assessment phases separated by a two-year interval. Before enrolment, participants confirmed that they had never consumed alcohol, had no family history of alcohol use disorders, and had no psychiatric or neurological conditions. They also completed the Alcohol Use Disorder Identification Test (AUDIT) (17) and participated in a semi-structured interview about their substance use habits. Those participants which reported consumption of any substance at this stage, we ruled out from the experiment. In the first phase—conducted before the onset of alcohol use—142 adolescents participated in the Magnetoencephalography (MEG) session, which involved five minutes of resting-state recording with eyes closed. Among them, 121 also underwent a magnetic resonance imaging (MRI) scan. After two years, 92 participants returned for a follow-up session, where they completed the AUDIT and semi-structured interview, and a similar MEG recording. Using consumption information, we calculated the number of Standard Alcohol Units (SAUs) typically consumed during drinking episodes, considering both the quantity and type of beverage consumed within a 2–3-hour period. To control for the effects of other substances, we excluded participants who reported regular tobacco or cannabis use. After quality control of MEG data from both sessions, 61 adolescents (35 males, 26 females; stage 1: mean age = 14.42 ± 0.64; stage 2: mean age = 16.40 ± 0.60) met all inclusion criteria and completed the full protocol. The Ethics Committee of the Universidad Complutense de Madrid approved the study, and all participants and their parents or legal guardians provided written informed consent in accordance with the Declaration of Helsinki.

### 2.2 MRI recordings

Participants underwent high-resolution 3D T1-weighted brain MRI scans using 1.5 Tesla machines at either the Santa Elena Foundation or the Clinical Hospital of Madrid. At the Santa Elena Foundation, imaging was conducted with a General Electric Optima MR450w scanner using the following parameters: echo time of 4.2 ms, repetition time of 11.2 ms, inversion time of 450 ms, flip angle of 12°, field of view of 100, acquisition matrix of 256 × 256, and a slice thickness of 1 mm. At the Clinical Hospital of Madrid, a General Electric Signa HDxt system was used with the same echo time, repetition time, and inversion time, but with a flip angle of 20°, while maintaining the same field of view, matrix size, and slice thickness.

### 2.3 MEG recordings and signal processing

MEG data were recorded with a 306-channel Elekta Neuromag system (102 magnetometers, 204 planar gradiometers) in a magnetically shielded room at the Center for Biomedical Technology, Madrid. Signals were band-pass filtered online between 0.1 and 330 Hz and sampled at 1,000 Hz. Head shapes were digitized using a 3D Fastrak system (Polhemus) by registering three fiducial points (nasion, left and right preauricular). Four Head Position Indicator (HPI) coils—two on the forehead and two on the mastoids—tracked head movements continuously during recording. Bipolar electrodes monitored eye movements (EOG) and cardiac activity (ECG) to identify physiological artifacts. To reduce environmental interference and correct for participant movement, the temporal extension of the Signal Space Separation (tSSS) method was applied, using a 10-second correlation window and a correlation threshold of 0.9 (18). Subsequently, the data were processed using the FieldTrip toolbox (19) in Matlab R2020b, where artifacts were automatically identified and subsequently verified by an experienced MEG analyst. Subsequently, EOG and ECG artifact removal were performed using an Independent Component Analysis (ICA) approach based on the Second Order Blind Identification (SOBI-3) algorithm, targeting components associated with ocular and cardiac activity. Eventually, clean segments of data were divided into 4-second epochs, with an additional 2 seconds of real signal added on either side as padding.

### 2.4 Source-space reconstruction

Source-level MEG signals were reconstructed using each participant’s individual T1-weighted MRI scan. A uniform grid of source locations, defined according to the Automated Anatomical Labeling (AAL) atlas (20), served as the basis for the source model. This grid included 1202 nodes corresponding to 78 cortical regions of interest, which were mapped into each subject’s anatomical space through a linear transformation from the MNI template to their personal MRI. The T1-weighted image was also used to generate a single-shell head model, derived from the inner skull surface (21). By integrating the head model, source model, and sensor layout, a leadfield matrix was computed using a modified spherical approach. For the source reconstruction, a linearly constrained minimum variance (LCMV) beamformer was applied as the inverse solution. To make magnetometer and gradiometer data compatible, the recordings and the leadfield matrix were normalized by channel, allowing for comparison of signal amplitudes across sensor types (22). Then, the three-dimensional time series at each source location was projected onto its main spatial principal component, resulting in a single time series per source point. To improve the accuracy of phase synchronization estimates, source-level activity was calculated separately for each traditional frequency band: theta (4–8 Hz), alpha (8–12 Hz), low beta (12–20 Hz), high beta (20–30 Hz), and low gamma (30–45 Hz). Each band was filtered using an FIR filter of order 1,800 with a Hamming window and implemented in a two-pass manner to mitigate any potential phase distortion.

### 2.5 Functional connectivity analysis

Functional connectivity (FC) was evaluated using the phase locking value (PLV), a metric that quantifies phase synchronization by analysing the phase differences between two time series. To compute the PLV across different frequency bands and time epochs, we followed the methodology outlined by Bruña et al., 2018. This approach produced symmetrical whole-brain connectivity matrices for each participant and frequency band, with dimensions of 1202[×[1202 nodes. The frequency bands analysed included theta (4–8 Hz), alpha (8–12 Hz), low-beta (12–20 Hz), high-beta (20–30 Hz), and gamma (30–45 Hz). For each pair of the 78 defined cortical regions, a PLV value was derived by calculating the root-mean-square of all PLV values linking nodes within those regions, resulting in a 78[×[78 FC matrix representing whole-brain connectivity.

### 2.6 Statistical analysis

#### Replication analysis during pre-consumption stage

The association between PLV values for each pair of ROIs and the Standard Alcohol Units (SAUs) was examined separately for each frequency band using a cluster-based permutation test (CBPT), following the approach described by Zalesky et al., 2010. To replicate the previous findings (13,25) in the longitudinal sample, one-tailed Spearman partial correlations, adjusted for age, sex, and data sample as covariates, were applied under the assumption of a positive relationship. Clusters were defined as groups of spatially contiguous connections showing significant partial correlations (with a p-value < 0.005), all in the same positive direction between PLV and SAU values. To ensure robustness, only clusters covering at least 1% of the source grid (i.e., involving a minimum of 12 nodes) were considered for further analysis. Spearman rho coefficients were transformed into Fisher Z scores, and a cluster-mass statistic was calculated by summing the Z values of all links within a given cluster. Statistical significance of each cluster was determined through a nonparametric procedure, using a null distribution generated from 50000 permutations of the dataset (randomly shuffling labels), as proposed by Maris & Oostenveld, 2007. Only clusters with a corrected p-value below 0.05 were retained for subsequent analyses. Given that electrophysiological recordings from EEG and MEG are often susceptible to volume conduction and signal leakage artifacts, all connections identified through the CBPT analysis were subsequently corrected by incorporating, as covariates, the correlation between the spatial filters derived from the beamformer solution and the spectral power within the frequency band of interest for the corresponding ROIs. This method is based on the idea that signal leakage is proportionally linked to the correlation between beamformer spatial filters of connected regions, and that higher activation power leads to a linear increase in leakage effects. Only those connections that remained statistically significant after this correction procedure were retained for further analysis. Finally, to evaluate the similarity of the networks extracted in the replication analysis, the Dice similarity coefficient (DSC) was computed using the networks reported in the original article as the gold standard:

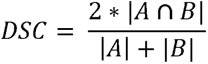

where ∣A∣ and ∣B∣ represent the number of connections in the original and replicated networks, respectively, and ∣A∩B∣ denotes the number of overlapping connections between them. The DSC ranges from 0 to 1, where 0 indicates no overlap and 1 indicates perfect similarity between the two networks.

#### Longitudinal analysis: General Linear Mixed Models

Functional connectivity values obtained in both stages of the networks identified in the CBPT analysis of the first phase were entered into a linear mixed-effects model (LMM). The model was specified to assess the effects of alcohol consumption and sex on the evolution of functional connectivity, with particular focus on the interaction effects between **consumption × stage** and **biological sex × stage**. The model was defined as:

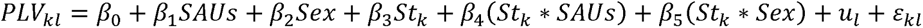

where *PLV_kl_* represents the functional connectivity value for stage *k* and subject *l*, *βs* are the fixed effects coefficients, *u_l_* denotes the random effect associated with the participant, and *ε_kl_* the residual error term. Additionally, the LMM was performed independently for each link of the network to provide a spatial localization of significant effects and individual statistical parameters. To account for multiple comparisons, the resulting p-values were corrected using the False Discovery Rate (FDR).

#### Functional Connectivity After Alcohol Intake

To assess the strength of the relationship between PLV and SAUs after alcohol intake, cluster-based permutation testing (CBPT) was performed using two-tailed Spearman correlations between PLV values from the second stage of the study and SAUs, restricting the analysis to the network clusters identified in the first stage. Clusters were defined as ensembles of spatially contiguous connection exhibiting statistically significant partial correlations (p < 0.05) and a uniform directionality of association across all constituent links. For each significant cluster, a representative PLV value was computed by averaging the scores within the cluster. Finally, the networks obtained in the CBPT analysis were corrected for the effects of volume conduction and leakage using the same methodology described in the replication analysis.

## 3. Results

During the second stage, adolescents displayed a range of drinking behaviors, from abstinence to heavy use, with a mean consumption of 3.96 ± 2.83 SAUs. To explore changes in functional connectivity associated with alcohol use, we first replicated the findings reported in our previous study (13), as the current cohort represents a subsample of that dataset. After confirming those results, we investigated factors influencing the longitudinal evolution of functional connectivity by testing interaction effects (predictor × stage) within linear mixed-effects models. Finally, we examined the functional connectivity patterns at follow-up as a function of current drinking levels.

### 3.1 Replication of FC-SAUs correlations

The correlations between PLV values and SAUs from the first stage were successfully replicated using a whole-brain CBPT analysis. Three of the five networks reported previously showed significant positive correlations: theta (4–8 Hz), alpha (8–12 Hz), and high beta (20–30 Hz). The theta network (Figure 1A) comprised 102 bilateral frontotemporal, parietal, and occipital connections (p < 0.05, ρ = 0.59); the alpha network (Figure 1B) included 91 predominantly left frontotemporal and parietal links (p < 0.05, ρ = 0.47); and the high-beta network (Figure 1C) consisted of 91 interhemispheric frontoparietal connections (p < 0.05, ρ = 0.55). After correcting volume conduction and leakage, all but two theta-band links remained significant. Similarity analyses showed moderate overlap in the theta band (Dice = 0.47) and higher reproducibility in the alpha (Dice = 0.64) and high-beta (Dice = 0.62) networks. A detailed view of the replicated and original connections is provided in Supplementary Material 1.

**Figure 1.**
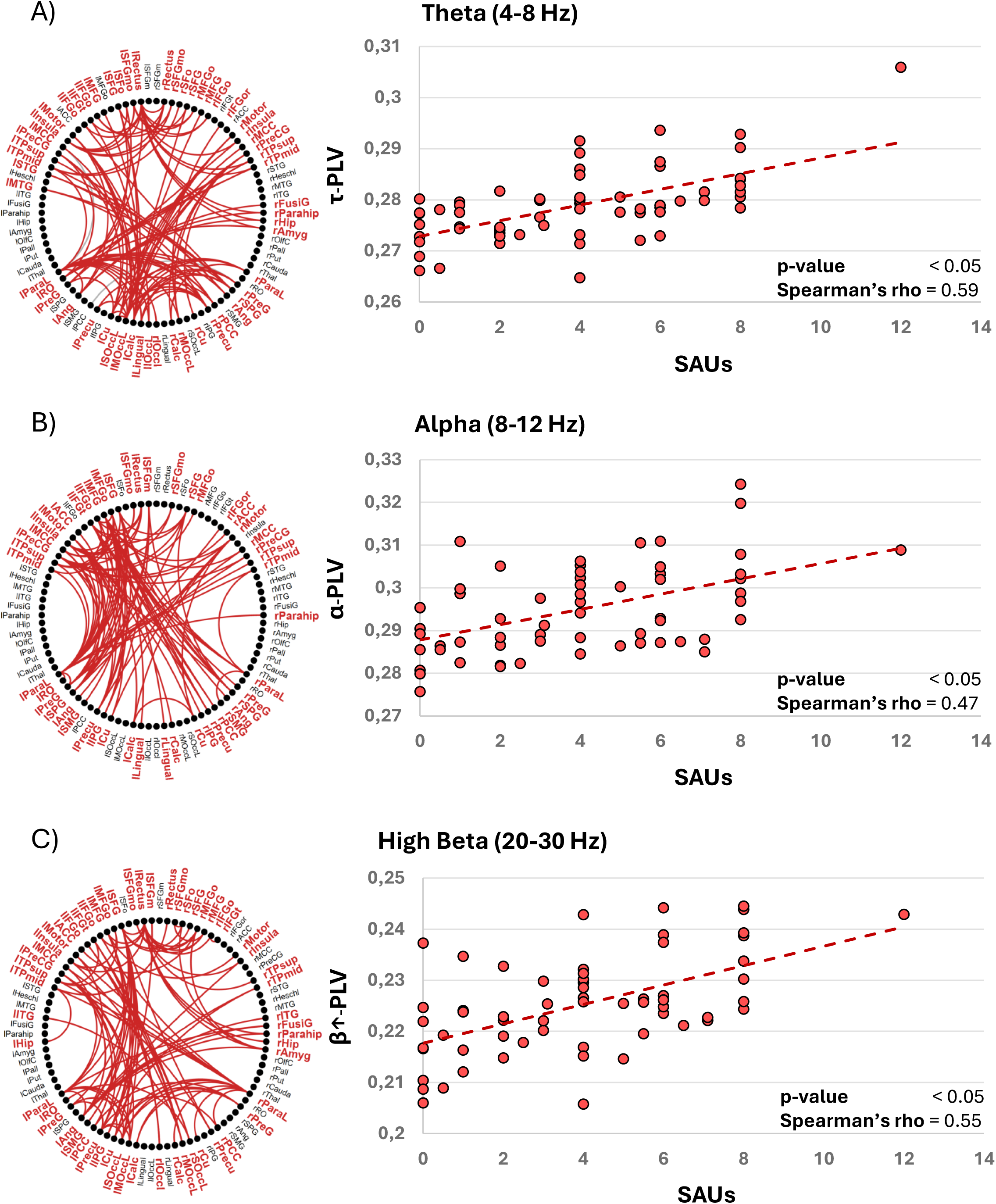
CBPT analysis of functional connectivity (PLV) with consumption levels (SAUs) during pre-consumption stage. On the left, the red dots represent the individual values and the red dashed line the trend line for the PLV-SAUs correlation. On the right, a summary graph of the spatial distribution of the network is shown. Grey lines represent those links that did not surpass the volume conduction and leakage correction. A) Correlation of theta PLV and SAUs. B) Correlation of alpha PLV and SAUs. C) Correlation of beta PLV and SAUs.

### 3.2 Key factors involved in the evolution of functional connectivity

Based on the functional networks extracted in the replication analysis, general linear mixed models were performed to determine the factors influencing functional connectivity throughout the study (Table 1). In the theta band the model revealed that study stage (*p* < 0.05, F = 5.22) and stage x SAUs interaction (*p* < 0.001, F = 11.86) were significant predictors of network PLV. Likewise, the individual link analysis showed that, of the 102 links comprising the network, 31 displayed a significant stage x SAUs interaction, of which only 8 survived FDR correction (FDR-threshold = 0.004) and were composed of frontal, temporoparietal, and occipital connections (Figure 2B). For the alpha band, only the stage x SAUs interaction (*p* < 0.01, F = 9.82) emerged as a significant predictor of network PLV. The individual link analysis identified a significant stage x SAUs interaction in 68 of the 91 links composing the network, 58 of which survived the threshold imposed by FDR correction (FDR-threshold = 0.03), distributed across the alpha network involving predominantly frontoparietal, and occipital regions (Figure 3B). Finally, in the high-beta band the model showed that both stage (*p* < 0.05, F = 4.94) and stage x SAUs interaction (*p* < 0.01, F = 10.04) acted as significant predictors of network PLV. Of the 91 models evaluated for each of the links comprising the network, 40 displayed a significant stage x SAUs interaction, of which 10 survived FDR correction (FDR-threshold = 0.002), distributed across connections mainly involving the left hemisphere in fronto-occipital, and temporal networks (Figure 4B). We plotted the mean PLV values for each stage and terciles groups based on SAU scores—Low Drinking (LD), Moderate Drinking (MD), and Binge Drinking (BD)— (Figure 2A, 3A, and 4A). The resulting patterns showed that LD and MD participants tended to increase their functional connectivity over time, whereas the BD group exhibited stable or decreasing PLV values across stages.

**Figure 2.**
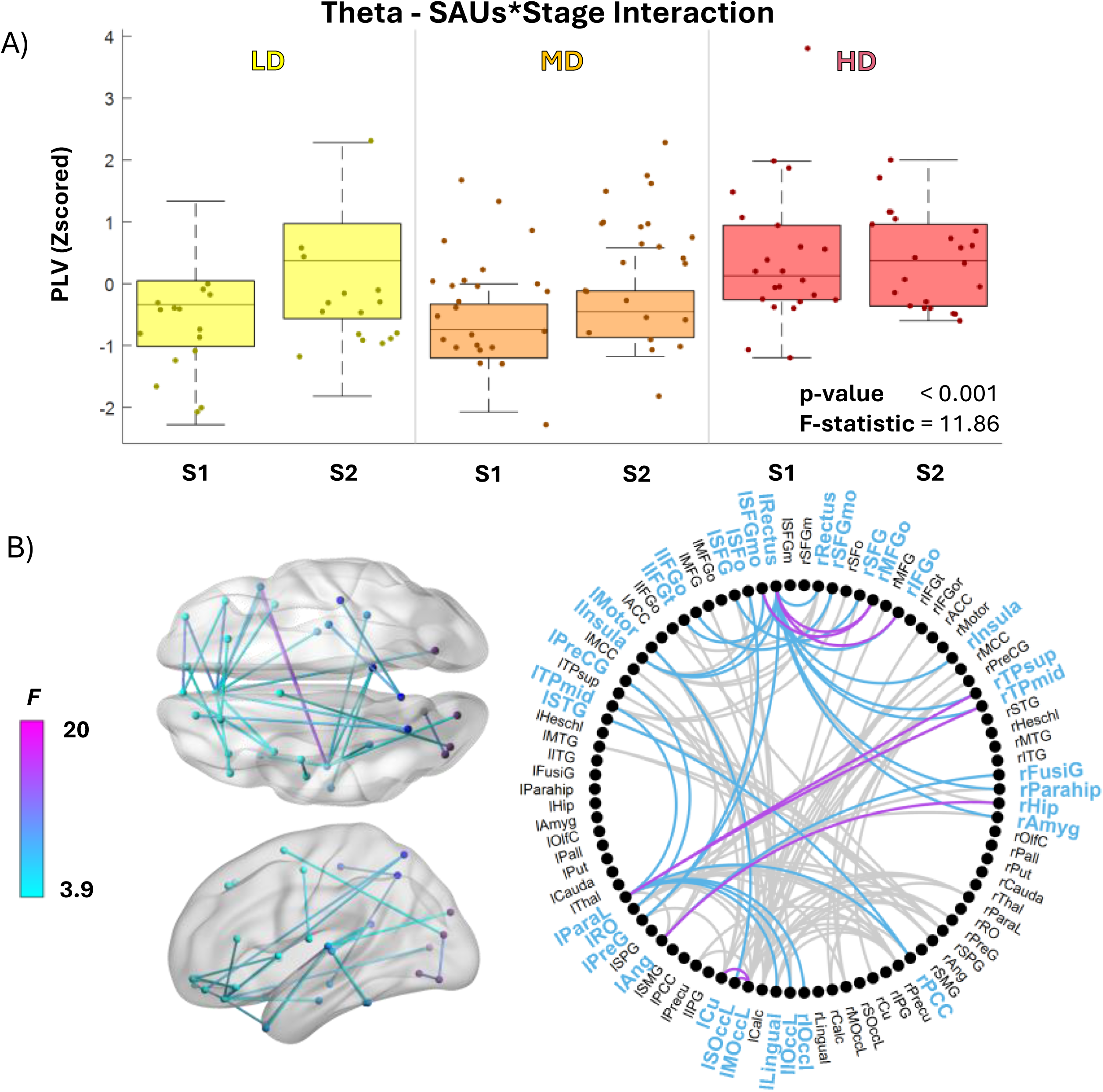
General linear mixed model of functional connectivity within the theta network in relation to alcohol consumption throughout the study. A) Functional connectivity within the theta network is represented for each stage of the study (S1 and S2) according to alcohol consumption pattern: light drinkers (LD = 0–1 SAUs) in yellow, moderate drinkers (MD = 2–5 SAUs) in orange and heavy drinkers (HD = 5.5–12 SAUs) in red. B) On the left, the distribution of links as a function of the statistic F on a template brain in horizontal and sagittal view is depicted. On the right there is a summary graph of the links with significant SAUs*Stage interaction. Purple lines represent the links that surpass FDR correction. Blue lines represent those links that did not surpass the FDR correction. Grey lines represent those links of the original theta network that did not show significant SAUs*Stage interaction.

**Figure 3.**
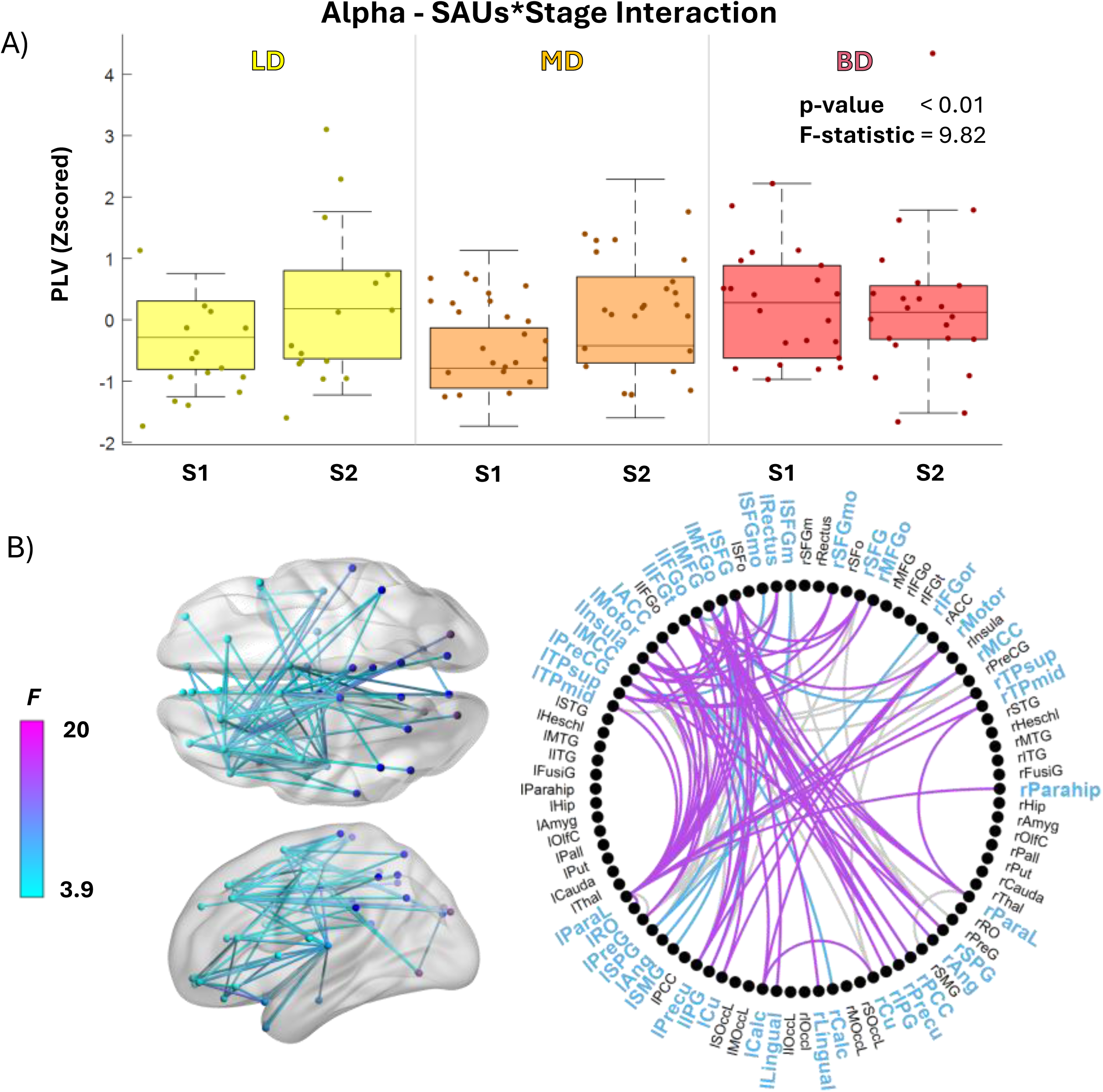
General linear mixed model of functional connectivity within the alpha network in relation to alcohol consumption throughout the study. A) Functional connectivity within the alpha network is represented for each stage of the study (S1 and S2) according to alcohol consumption pattern: light drinkers (LD = 0–1 SAUs) in yellow, moderate drinkers (MD = 2–5 SAUs) in orange and heavy drinkers (HD = 5.5–12 SAUs) in red. B) On the left, the distribution of links as a function of the statistic F on a template brain in horizontal and sagittal view is depicted. On the right there is a summary graph of the links with significant SAUs*Stage interaction. Purple lines represent the links that surpass FDR correction. Blue lines represent those links that did not surpass the FDR correction. Grey lines represent those links of the original alpha network that did not show significant SAUs*Stage interaction.

**Figure 4.**
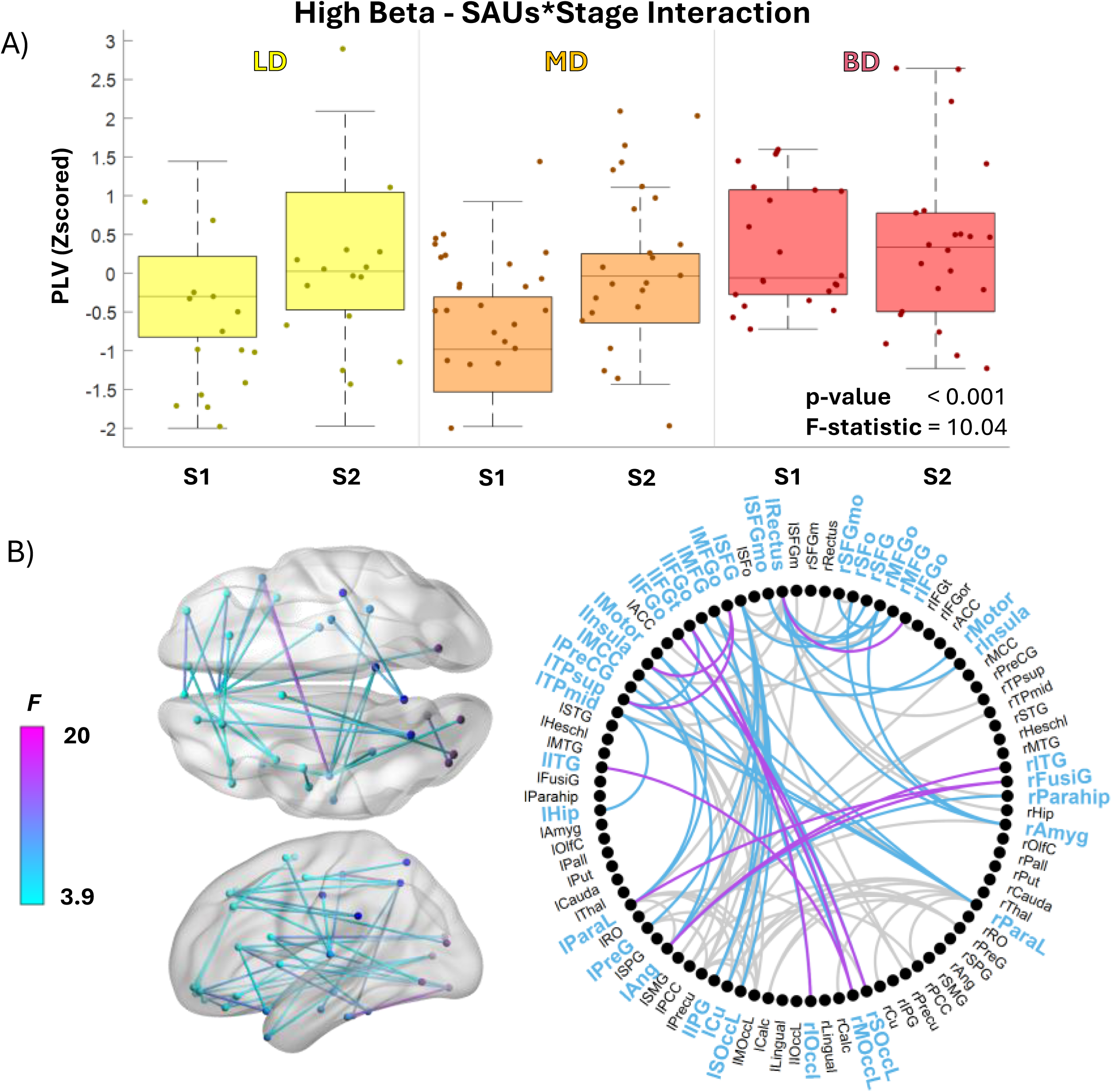
General linear mixed model of functional connectivity within the high beta network in relation to alcohol consumption throughout the study. A) Functional connectivity within the high beta network is represented for each stage of the study (S1 and S2) according to alcohol consumption pattern: light drinkers (LD = 0–1 SAUs) in yellow, moderate drinkers (MD = 2–5 SAUs) in orange and heavy drinkers (HD = 5.5–12 SAUs) in red. B) On the left, the distribution of links as a function of the statistic F on a template brain in horizontal and sagittal view is depicted. On the right there is a summary graph of the links with significant SAUs*Stage interaction. Purple lines represent the links that surpass FDR correction. Blue lines represent those links that did not surpass the FDR correction. Grey lines represent those links of the original high beta network that did not show significant SAUs*Stage interaction.

**Table 1.** General Linear Mixed Model (GLMM) analyses to predict fuctional connectivity (PLV) using stage, alcohol consumption (SAUs), sex, age difference and stage interactions as predictors.

| **N = 61** | **F-statistic** | **p-val** |
| --- | --- | --- |
| **Theta (4-8 Hz)** |  |  |
| *Intercept* | 1.00 | 0.32 |
| **Stage** | **5.22** | **< 0.05\*** |
| SAUs | 3.73 | 0.06 |
| Sex | 0.01 | 0.93 |
| Age difference (S2-S1) | 0.12 | 0.72 |
| **Stage \* SAUs Interaction** | **11.86** | **< 0.001\*** |
| Stage * Sex Interaction | 0.19 | 0.66 |
| Stage * Age difference Interaction | 0.86 | 0.35 |
| **Alpha (8-12 Hz)** |  |  |
| *Intercept* | 1.10 | 0.30 |
| Stage | 0.77 | 0.38 |
| SAUs | 0 | 0.99 |
| Sex | 0.14 | 0.71 |
| Age difference (S2-S1) | 0.41 | 0.52 |
| **Stage \* SAUs Interaction** | **9.82** | **< 0.01\*** |
| Stage * Sex Interaction | 1.02 | 0.31 |
| Stage * Age difference Interaction | 0.37 | 0.54 |
| **High beta (20-30 Hz)** |  |  |
| *Intercept* | 1.23 | 0.27 |
| **Stage** | **4.94** | **< 0.05\*** |
| SAUs | 0.46 | 0.50 |
| Sex | 0.05 | 0.82 |
| Age difference (S2-S1) | 0.05 | 0.82 |
| **Stage \* SAUs Interaction** | **10.04** | **< 0.01\*** |
| Stage * Sex Interaction | 0.12 | 0.73 |
| Stage * Age difference Interaction | 0.42 | 0.52 |

Although none of the networks showed significant main effects of sex or stage by sex interactions, analyses at the individual connection level revealed several significant links that did not survive FDR correction. In the theta band, 22 connections showed a stage × sex interaction, 10 of which overlapped with those also exhibiting a stage × SAUs interaction. In the alpha band, 10 connections showed this interaction, with 7 overlapping links, and in the high-beta band, 8 connections were significant, with 5 overlapping. The spatial distribution of these effects is presented in Supplementary Material 2.

### 3.3 FC-SAUs correlation after consumption

To describe the relationship between functional connectivity and alcohol consumption after drinking onset, an exploratory analysis was performed within the networks identified in the replication analysis. Following the changes observed throughout the study, the network described in the theta band showed a significant relationship between alcohol consumption and functional connectivity (*p* < 0.001, ρ = 0.54). The cluster formed within this network (Figure 5B) consisted of 38 links connecting occipital and parietal regions with bilateral frontal regions, 31 of which remained significant after correction for volume conduction and leakage effects. Regarding the networks described in the alpha and high-beta bands, no significant relationships were found between functional connectivity and alcohol consumption during the post-onset stage of the study.

**Figure 5.**
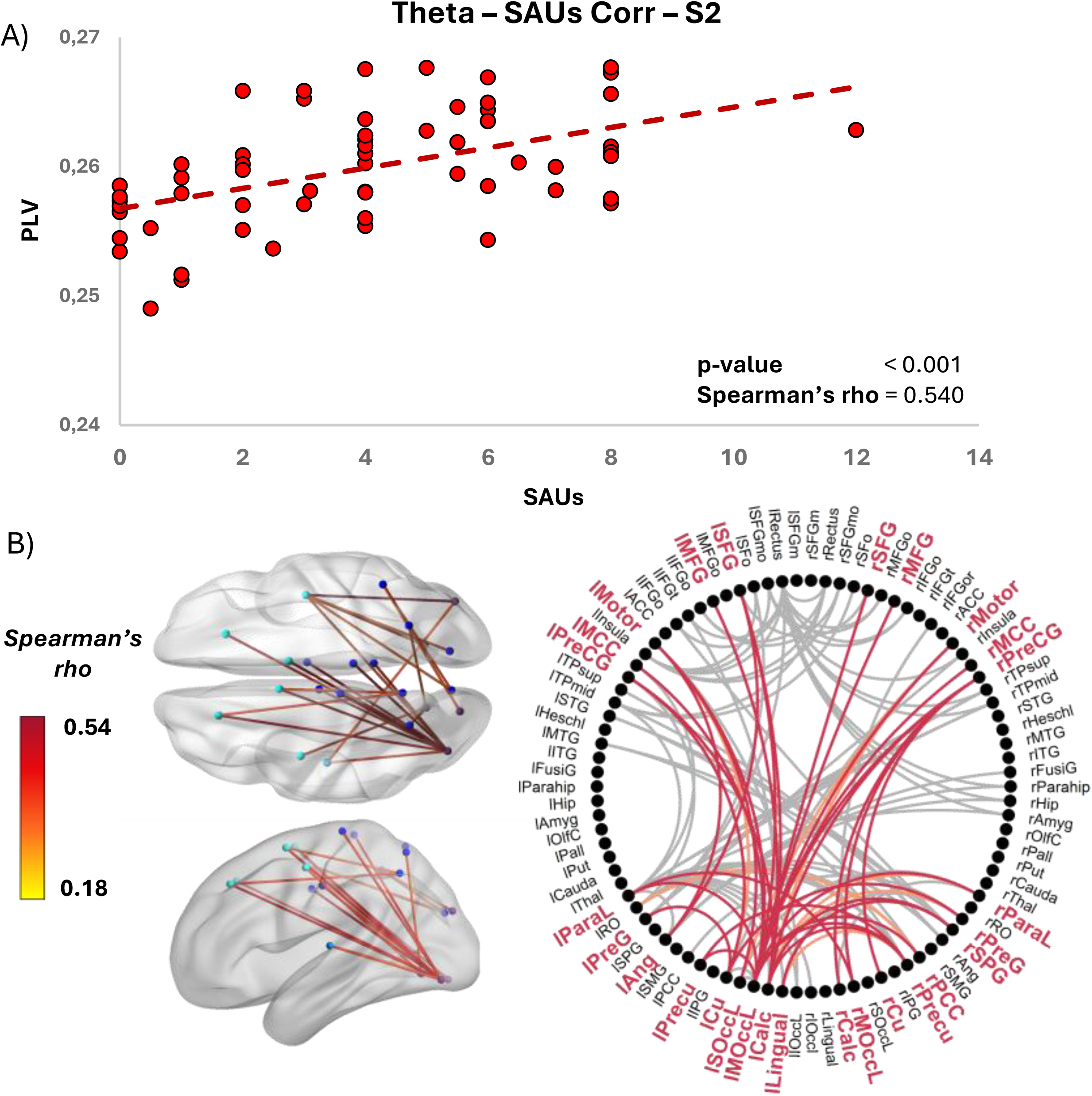
CBPT analysis of theta functional connectivity (PLV) with consumption levels (SAUs) during post-consumption stage. A) Correlation of theta PLV with SAUs. Red dots represent the individual values and the red dashed line the trend line for the PLV-SAUs correlation. B) On the left, the distribution of links as a function of the Spearman’s rho on a template brain in horizontal and sagittal view is depicted. On the right there is a summary graph of the links with significant theta PLV-SAUs correlation. Red lines represent the links within the cluster that surpass the volume conduction and leakage correction. Orange lines represent those links within the cluster that did not surpass the volume conduction and leakage correction. Grey lines represent those links of the original theta network out of the cluster.

## 4. Discussion

This study investigated longitudinal changes in functional connectivity (FC) patterns following the onset of alcohol consumption during early adolescence. The main findings revealed pre-existing FC differences that evolved distinctively according to the severity of subsequent alcohol use. Adolescents who engaged in heavier drinking showed a stabilization or reduction of FC within theta, alpha, and beta bands, contrasting with the steady increase observed in those with moderate or low alcohol intake. Furthermore, after the onset of alcohol misuse, higher anteroposterior theta-band connectivity was robustly associated with greater alcohol consumption.

### 4.1 Variability during neurodevelopmental synchronization

During adolescence, cortical remodeling unfolds through dynamic changes in dendritic arborization, microglial activity, myelination (2), GABAergic signaling (27), and dopamine levels (3). These neurodevelopmental processes reshape the balance between excitation and inhibition and are thought to support progressive increases in functional connectivity, reflected in enhanced network coherence, reduced variability, shorter communication paths to hub regions, and an improved capacity to recruit task-relevant nodes (28,29). Consistent with this framework, previous electrophysiological studies have characterized adolescence as a period of increasing oscillatory coupling and large-scale network integration (30–33), processes that enhance functional integration while gradually constraining plasticity and promoting network stabilization (34–36). Importantly, these maturational processes do not unfold uniformly across individuals. The dual systems model of neurodevelopment posits an imbalance between the relatively early maturation of subcortical reward circuits and the slower development of cortical regulatory systems (3). Moreover, cortical maturation follows network-specific trajectories, with sensory systems stabilizing earlier and association areas exhibiting more heterogeneous and prolonged development (35,36). Such differentiation likely shapes distinct cognitive and behavioral trajectories (36) and may heighten some adolescents’ vulnerability to earlier and more hazardous substance use. Our findings align with this perspective by revealing variability in neuromaturational profiles across adolescents. Specifically, individuals who reported low or moderate alcohol consumption exhibited lower functional connectivity prior to alcohol use onset, followed by the typical longitudinal increase. In contrast, adolescents with heavier alcohol consumption showed higher pre-consumption functional connectivity, followed by stabilization or even a reduction after alcohol use onset. Together, these results suggest a heterogeneous developmental trajectory of oscillatory synchronization, with heavier drinking patterns associated with deviations from the typical maturational course.

Within a broader neurodevelopmental framework, sex-related differences have been proposed, with several studies reporting earlier maturation in females, particularly between 14 and 16 years of age (37). However, findings in younger adolescents (9–12 years) have been less consistent (38). In the present study, although sex was examined as a potential moderator of neurodevelopmental trajectories and functional connectivity patterns, the observed effects were not sufficiently robust to support reliable sex-related differences within this age range.

### 4.2 Early stabilization of beta and alpha networks at rest

One possible interpretation of the different maturation trajectories observed in this study is that subjects with higher alcohol consumption exhibit a premature stabilization of cortical networks, consistent with a state of “pseudomaturation” previously linked to increased vulnerability to risky behaviors (11,39,40). Within this framework, elevated functional connectivity may reflect an apparently advanced neurodevelopmental stage, potentially driven by an accelerated developmental tempo shaped by interindividual differences in pubertal timing and substantial hereditary influences (37,38,41,42). This account is consistent with our findings showing an earlier increase in long-distance connectivity between prefrontal, parietal, and occipital regions, followed by a progressive convergence of alpha and beta connectivity by late adolescence. Resting-state alpha rhythms are known to continue maturing throughout adolescence (43,44) and support large-scale posterior–anterior integration, with parieto–occipital regions functioning as major hubs projecting toward frontal areas. Such alpha-mediated connectivity has been linked to the regulation of cortical excitability and the gating of distributed information across networks (45). By contrast, beta-band connectivity is typically more spatially focal, primarily engaging sensorimotor and parietal–frontal circuits, and is thought to sustain the current cognitive and sensorimotor state, thereby promoting network stability rather than flexible updating (45–47). Taken together, these frequency-specific patterns suggest that the observed trajectories may reflect an early shift from broadly integrative dynamics toward more stabilized and functionally specialized network configurations, consistent with a pseudomaturational profile in adolescents with heavier alcohol use.

### 4.3 Theta hyperconnectivity after alcohol use onset

During neurodevelopment, theta-band connectivity has been implicated in top-down coordination and the regulation of cortical excitability, facilitating efficient global information transfer (33). Nevertheless, it should be noted that, unlike the networks described in alpha and beta bands, FC in theta network remained hyperconnected in heavy drinkers after follow-up period. Consistent with this pattern, studies examining problematic alcohol use in university populations have reported increased theta connectivity across posterior interhemispheric and fronto-parietal networks (48–51). Given that alcohol consumption in those samples typically began during adolescence, some authors have suggested that such alterations may reflect alcohol-related interference with the maturation of large-scale brain networks. Within this framework, excess theta connectivity has been interpreted as a compensatory response in a brain affected by repeated alcohol exposure (52,53). However, our findings indicate that theta-band differences were already detectable prior to alcohol use initiation. This suggests that theta hyperconnectivity may not solely represent a consequence of alcohol exposure, but rather a pre-existing vulnerability marker that is subsequently sustained or amplified by heavy drinking, particularly within posterior regions of the default mode network (DMN). Supporting this interpretation, findings from the Collaborative Study of the Genetics of Alcoholism (COGA) show that polygenic variants associated with alcohol use disorder risk are also linked to increased theta connectivity (54), reinforcing the notion that this electrophysiological profile may index inherited susceptibility.

### 4.4 Strengths and limitations

Among the strengths of this study is its longitudinal design, which allows for the examination of functional network evolution across neurodevelopment and enables the dissociation of predispositional traits from potential effects of alcohol consumption. In addition, the combination of magnetoencephalography (MEG) and MRI provides measures of brain electrical activity with excellent temporal resolution and relatively high spatial resolution. Regarding limitations, MEG recordings cannot reliably reconstruct subcortical activity, which also plays a critical role in neurodevelopment and substance use. Moreover, part of the existing evidence on functional connectivity during adolescence and in the context of substance use is derived from fMRI data, which limits direct comparability across studies. Nevertheless, there is evidence supporting a high degree of correspondence between fMRI-based connectivity metrics and M/EEG measures.

### 4.5 Conclusion

In conclusion, our results reveal distinct neuromaturational trajectories associated with alcohol consumption, characterized by an early stabilization of functional networks in individuals with higher alcohol use and continued network development in moderate drinkers and abstinent adolescents. These differences may be explained by asynchronous neurodevelopmental processes that progressively converge across adolescence as individuals reach comparable levels of cortical maturation. Notably, beyond a limited set of pre-existing neural traits, such as theta-band hyperconnectivity, we did not observe clear effects of alcohol consumption, potentially reflecting a relatively short drinking history in this sample. Taken together, these findings support the view of neurodevelopment as a highly individualized process and highlight the importance of considering interindividual variability when addressing the rising prevalence of alcohol-related problems during adolescence and the risks associated with this critical developmental period.

## Supporting information

SuppMaterial

## 5. Aknowledgments

This work was supported by grants from the **Plan Nacional sobre Drogas (PNSD), Ministerio de Sanidad (MISAN)** (2021|075 to F.M.; 2017I039 and 2014I035 to F.M. and L.M.G.M.) and by the **Fundación Banco Santander (FBS)** (CT58/21–CT59/21 to A.d.C.-L.; CT15/23 to D.S.-B. and M.U.). The authors gratefully acknowledge this financial support.

**Alberto del Cerro-León:** Investigation; Formal analysis; Writing – original draft.

**Danylyna Shpakivska-Bilan:** Investigation.

**Marcos Uceta:** Investigation.

**Fernando Maestú:** Conceptualization; Funding acquisition.

**Luis Miguel García-Moreno:** Conceptualization; Funding acquisition.

**Luis Fernando Antón-Toro:** Conceptualization; Investigation; Writing – original draft.

## 7. Data availability

Processed data and scripts for statistics are publicly available on the OSF website (https://osf.io/276zy/). Raw data can be accessed through a data transfer agreement with the responsible university (Universidad Complutense de Madrid). Finally, the MEG signal preprocessing and cleaning codes can be found publicly available on the Github repository (https://github.com/rbruna/meeg_analysis).

---

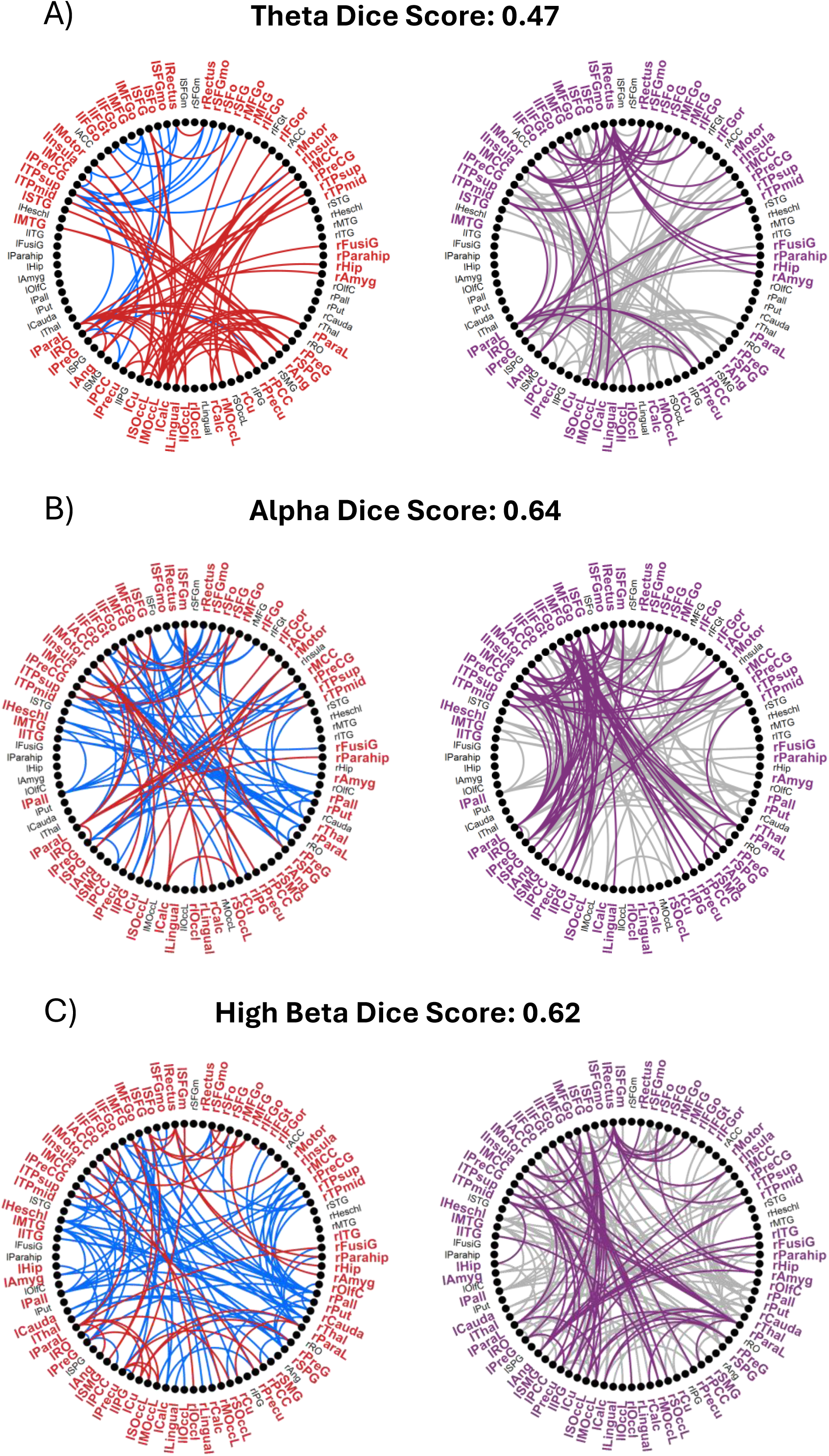

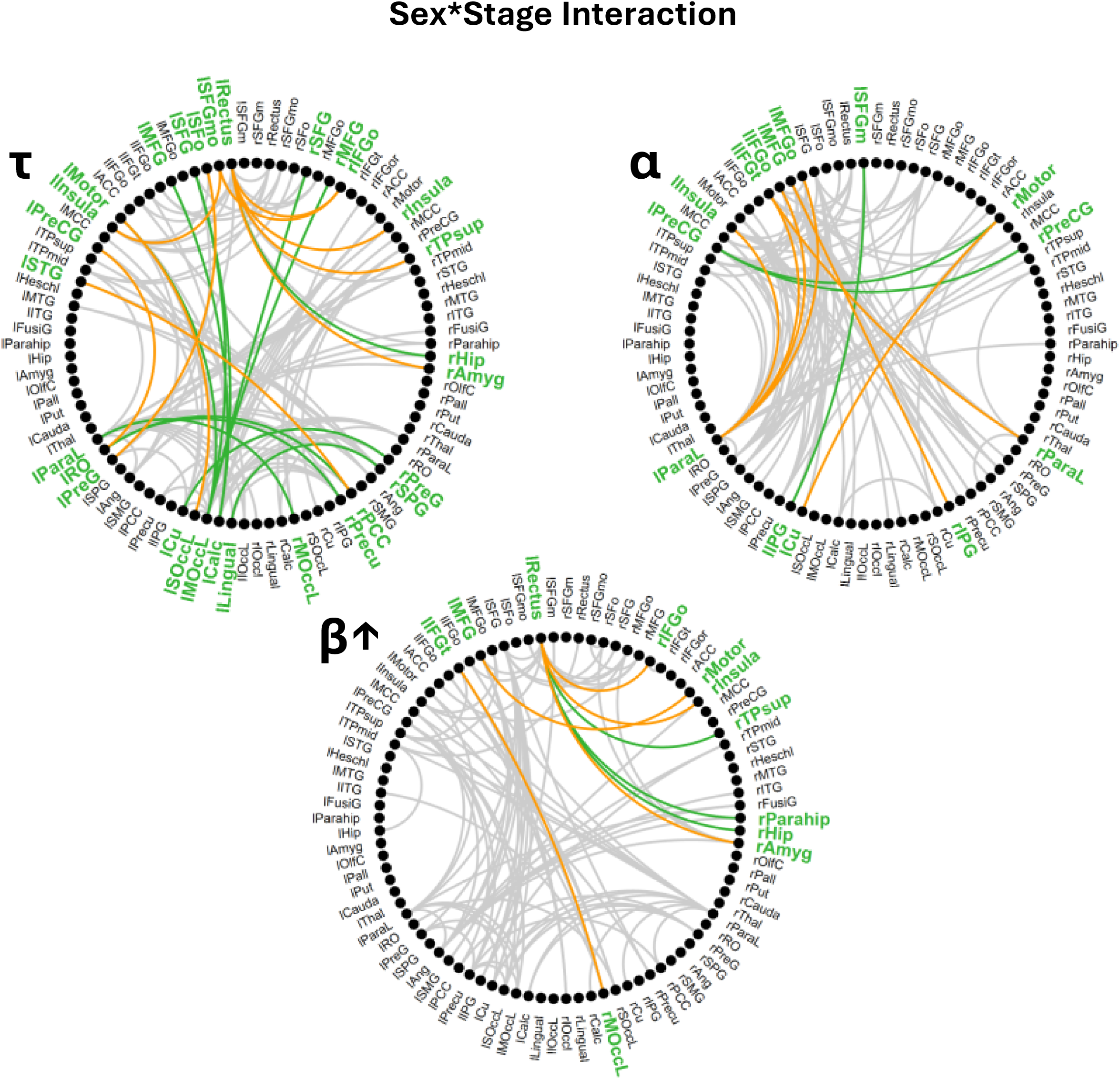

## References

1. Casey BJ, Jones RM, Hare TA (2008): The Adolescent Brain. Ann N Y Acad Sci 1124: 111–126.

2. Blakemore S-J, Choudhury S (2006): Development of the adolescent brain: implications for executive function and social cognition. J Child Psychol Psychiatry 47: 296–312.

3. Shulman EP, Smith AR, Silva K, Icenogle G, Duell N, Chein J, Steinberg L (2016): The dual systems model: Review, reappraisal, and reaffirmation. Dev Cogn Neurosci 17: 103–117.

4. Bava S, Tapert SF (2010): Adolescent brain development and the risk for alcohol and other drug problems. Neuropsychol Rev 20: 398–413.

5. Lees B, Debenham J, Squeglia LM (2021): Alcohol and Cannabis Use and the Developing Brain. Alcohol Res 41: 11.

6. Hammer JH, Parent MC, Spiker DA, World Health Organization (2018): Global status report on alcohol and health 2018. Global Status Report on Alcohol, vol. 65. 10.1037/cou0000248

7. Pfefferbaum A, Kwon D, Brumback T, Thompson WK, Cummins K, Tapert SF, et al. (2018): Altered Brain Developmental Trajectories in Adolescents After Initiating Drinking. Am J Psychiatry 175: 370–380.

8. Pérez-García JM, Suárez-Suárez S, Doallo S, Cadaveira F (2022): Effects of binge drinking during adolescence and emerging adulthood on the brain: A systematic review of neuroimaging studies. Neurosci Biobehav Rev 137: 104637.

9. Almeida-Antunes N, Crego A, Carbia C, Sousa SS, Rodrigues R, Sampaio A, López-Caneda E (2021): Electroencephalographic signatures of the binge drinking pattern during adolescence and young adulthood: A PRISMA-driven systematic review. NeuroImage Clin 29. 10.1016/j.nicl.2020.102537

10. Kühn S, Mascharek A, Banaschewski T, Bodke A, Bromberg U, Büchel C, et al. (2019): Predicting development of adolescent drinking behaviour from whole brain structure at 14 years of age. Elife 8: 1–15.

11. Squeglia LM, Ball TM, Jacobus J, Brumback T, McKenna BS, Nguyen-Louie TT, et al. (2017): Neural predictors of initiating alcohol use during adolescence. Am J Psychiatry 174: 172–185.

12. del Cerro-León A, Uceta M, Shpakivska-Bilan D, Suárez-Méndez I, Peribáñez-Baz H, Cuesta P, et al. (2025): Electrophysiological sexual dimorphism as an early risk marker of alcohol use in adolescence: A longitudinal neuroimaging study. Addiction 1–11.

13. del Cerro-León A, Fernando Antón-Toro L, Shpakivska-Bilan D, Uceta M, Santos-Mayo A, Cuesta P, et al. (2024): Adolescent alcohol consumption predicted by differences in electrophysiological functional connectivity and neuroanatomy. Proc Natl Acad Sci 121. 10.1073/pnas.2320805121

14. Antón-Toro LF, Bruña R, Suárez-Méndez I, Correas Á, García-Moreno LM, Maestú F (2021): Abnormal organization of inhibitory control functional networks in future binge drinkers. Drug Alcohol Depend 218. 10.1016/j.drugalcdep.2020.108401

15. Antón-Toro LF, Shpakivska-Bilan D, López-Abad L, Del Cerro-León A, Uceta M, Bruña R, et al. (2026): Adolescent predisposition to binge drinking is associated with differences in inhibitory control MEG event-related fields. Front Psychiatry 16. 10.3389/fpsyt.2025.1696748

16. Ruan H, Zhou Y, Luo Q, Robert GH, Desrivières S, Quinlan EB, et al. (2019): Adolescent binge drinking disrupts normal trajectories of brain functional organization and personality maturation. NeuroImage Clin 22: 101804.

17. Guillamón C, Solé G, Farran C (1999): Test para la identificación de transtornos por uso de alcohol (AUDIT): Traducción y validación del AUDIT al catalán y castellano. Adicciones 11: 347.

18. Taulu S, Simola J (2006): Spatiotemporal signal space separation method for rejecting nearby interference in MEG measurements. Phys Med Biol 51: 1759–1768.

19. Oostenveld R, Fries P, Maris E, Schoffelen JM (2011): FieldTrip: Open source software for advanced analysis of MEG, EEG, and invasive electrophysiological data. Comput Intell Neurosci 2011. 10.1155/2011/156869

20. Tzourio-Mazoyer N, Landeau B, Papathanassiou D, Crivello F, Etard O, Delcroix N, et al. (2002): Automated anatomical labeling of activations in SPM using a macroscopic anatomical parcellation of the MNI MRI single-subject brain. Neuroimage 15: 273–289.

21. Nolte G (2003): The magnetic lead field theorem in the quasi-static approximation and its use for magnetoenchephalography forward calculation in realistic volume conductors. Phys Med Biol 48: 3637–3652.

22. Mohseni HR, Woolrich MW, Kringelbach ML, Luckhoo H, Smith PP, Aziz TZ (2012): Fusion of Magnetometer and Gradiometer Sensors of MEG in the Presence of Multiplicative Error. IEEE Trans Biomed Eng 59: 1951–1961.

23. Bruña R, Maestú F, Pereda E (2018): Phase locking value revisited: Teaching new tricks to an old dog. J Neural Eng 15. 10.1088/1741-2552/aacfe4

24. Zalesky A, Fornito A, Bullmore ET (2010): Network-based statistic: Identifying differences in brain networks. Neuroimage 53: 1197–1207.

25. Antón-Toro LF, Bruña R, Del Cerro-León A, Shpakivska D, Mateos-Gordo P, Porras-Truque C, et al. (2022): Electrophysiological resting-state hyperconnectivity and poorer behavioural regulation as predisposing profiles of adolescent binge drinking. Addict Biol 27: 1–11.

26. Maris E, Oostenveld R (2007): Nonparametric statistical testing of EEG- and MEG-data. J Neurosci Methods 164: 177–190.

27. Caballero A, Flores-Barrera E, Cass DK, Tseng KY (2014): Differential regulation of parvalbumin and calretinin interneurons in the prefrontal cortex during adolescence. Brain Struct Funct 219: 395–406.

28. Lockhart S, Sawa A, Niwa M (2018): Developmental trajectories of brain maturation and behavior: Relevance to major mental illnesses. J Pharmacol Sci 137: 1–4.

29. Meng L, Xiang J (2016): Frequency specific patterns of resting-state networks development from childhood to adolescence: A magnetoencephalography study. Brain Dev 38: 893–902.

30. Schäfer CB, Morgan BR, Ye AX, Taylor MJ, Doesburg SM (2014): Oscillations, networks, and their development: MEG connectivity changes with age. Hum Brain Mapp 35: 5249–5261.

31. Briley PM, Liddle EB, Groom MJ, Smith HJF, Morris PG, Colclough GL, et al. (2018): Development of human electrophysiological brain networks. J Neurophysiol 120: 3122–3130.

32. Brookes MJ, Groom MJ, Liuzzi L, Hill RM, Smith HJF, Briley PM, et al. (2018): Altered temporal stability in dynamic neural networks underlies connectivity changes in neurodevelopment. Neuroimage 174: 563–575.

33. Hunt BAE, Wong SM, Vandewouw MM, Brookes MJ, Dunkley BT, Taylor MJ (2019): Spatial and spectral trajectories in typical neurodevelopment from childhood to middle age. Netw Neurosci 3: 497–520.

34. Glasser MF, Goyal MS, Preuss TM, Raichle ME, Van Essen DC (2014): Trends and properties of human cerebral cortex: Correlations with cortical myelin content. Neuroimage 93: 165–175.

35. Hettwer MD, Dorfschmidt L, Puhlmann LMC, Jacob LM, Paquola C, Bethlehem RAI, et al. (2024): Longitudinal variation in resilient psychosocial functioning is associated with ongoing cortical myelination and functional reorganization during adolescence. Nat Commun 15. 10.1038/s41467-024-50292-2

36. Park BY, Paquola C, Bethlehem RAI, Benkarim O, Mišić B, Smallwood J, et al. (2022): Adolescent development of multiscale structural wiring and functional interactions in the human connectome. Proc Natl Acad Sci U S A 119: 1–11.

37. Brouwer RM, Schutte J, Janssen R, Boomsma DI, Hulshoff Pol HE, Schnack HG (2021): The Speed of Development of Adolescent Brain Age Depends on Sex and Is Genetically Determined. Cereb Cortex 31: 1296–1306.

38. Holm MC, Leonardsen EH, Beck D, Dahl A, Kjelkenes R, de Lange AMG, Westlye LT (2023): Linking brain maturation and puberty during early adolescence using longitudinal brain age prediction in the ABCD cohort. Dev Cogn Neurosci 60: 101220.

39. Berns GS, Moore S, Capra CM (2009): Adolescent engagement in dangerous behaviors is associated with increased white matter maturity of frontal cortex. PLoS One 4. 10.1371/journal.pone.0006773

40. Kardan O, Weigard AS, Cope LM, Martz ME, Angstadt M, McCurry KL, et al. (2025): Functional Brain Connectivity Predictors of Prospective Substance Use Initiation and Their Environmental Correlates. Biol Psychiatry Cogn Neurosci Neuroimaging 10: 203–212.

41. Bottenhorn KL, Cardenas-Iniguez C, Mills KL, Laird AR, Herting MM (2023): Profiling intra- and inter-individual differences in brain development across early adolescence. Neuroimage 279: 120287.

42. Mccann CF, Cheng TW, Mills KL, Silvers JA (2026): Developmental Cognitive Neuroscience Pubertal timing and tempo differentially influence cortical and subcortical maturation in adolescence. Dev Cogn Neurosci 77: 101657.

43. Cellier D, Riddle J, Petersen I, Hwang K (2021): The development of theta and alpha neural oscillations from ages 3 to 24 years. Dev Cogn Neurosci 50: 100969.

44. Rubia K, Smith AB, Woolley J, Nosarti C, Heyman I, Taylor E, Brammer M (2006): Progressive increase of frontostriatal brain activation from childhood to adulthood during event[related tasks of cognitive control. Hum Brain Mapp 27: 973–993.

45. Hillebrand A, Tewarie P, van Dellen E, Yu M, Carbo EWS, Douw L, et al. (2016): Direction of information flow in large-scale resting-state networks is frequency-dependent. Proc Natl Acad Sci 113: 3867–3872.

46. Engel AK, Fries P (2010): Beta-band oscillations — signalling the status quo? Curr Opin Neurobiol 20: 156–165.

47. Brookes MJ, Woolrich M, Luckhoo H, Price D, Hale JR, Stephenson MC, et al. (2011): Investigating the electrophysiological basis of resting state networks using magnetoencephalography. Proc Natl Acad Sci 108: 16783–16788.

48. Jansen JM, van Wingen G, van den Brink W, Goudriaan AE (2015): Resting state connectivity in alcohol dependent patients and the effect of repetitive transcranial magnetic stimulation. Eur Neuropsychopharmacol 25: 2230–2239.

49. Lesnewich LM, Pawlak AP, Gohel S, Bates ME (2022): Functional connectivity in the central executive network predicts changes in binge drinking behavior during emerging adulthood: an observational prospective study. Addiction 117: 1899–1907.

50. Zhu X, Cortes CR, Mathur K, Tomasi D, Momenan R (2017): Model[free functional connectivity and impulsivity correlates of alcohol dependence: a resting[state study. Addict Biol 22: 206–217.

51. Correas A, Rodriguez Holguín S, Cuesta P, López-Caneda E, García-Moreno LM, Cadaveira F, Maestú F (2015): Exploratory Analysis of Power Spectrum and Functional Connectivity During Resting State in Young Binge Drinkers: A MEG Study. Int J Neural Syst 25: 1550008.

52. Kim BM, Kim MS, Kim JS (2020): Alterations of Functional Connectivity During the Resting State and Their Associations With Visual Memory in College Students Who Binge Drink. Front Hum Neurosci 14: 1–11.

53. Correas A, Cuesta P, López-Caneda E, Rodríguez Holguín S, García-Moreno LM, Pineda-Pardo JA, et al. (2016): Functional and structural brain connectivity of young binge drinkers: A follow-up study. Sci Rep 6. 10.1038/srep31293

54. Meyers JL, Chorlian DB, Johnson EC, Pandey AK, Kamarajan C, Salvatore JE, et al. (2019): Association of polygenic liability for alcohol dependence and EEG connectivity in adolescence and young adulthood. Brain Sci 9: 1–17.

