## Supplementary material for "Neurodevelopment Trajectory of Electrophysiological Functional Connectivity Following Alcohol Use Initiation": SuppMaterial

**Supplementary Material 1: Replication of FC-SAUs correlations**

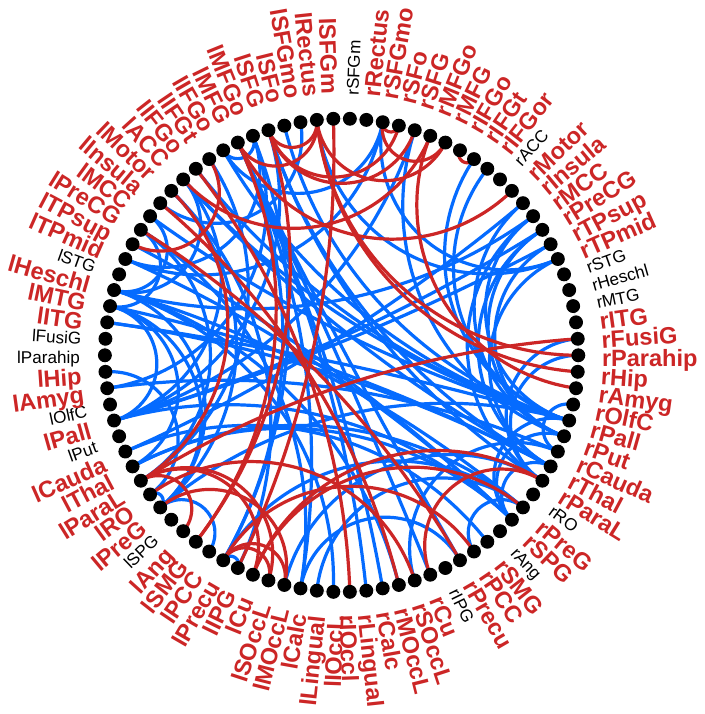

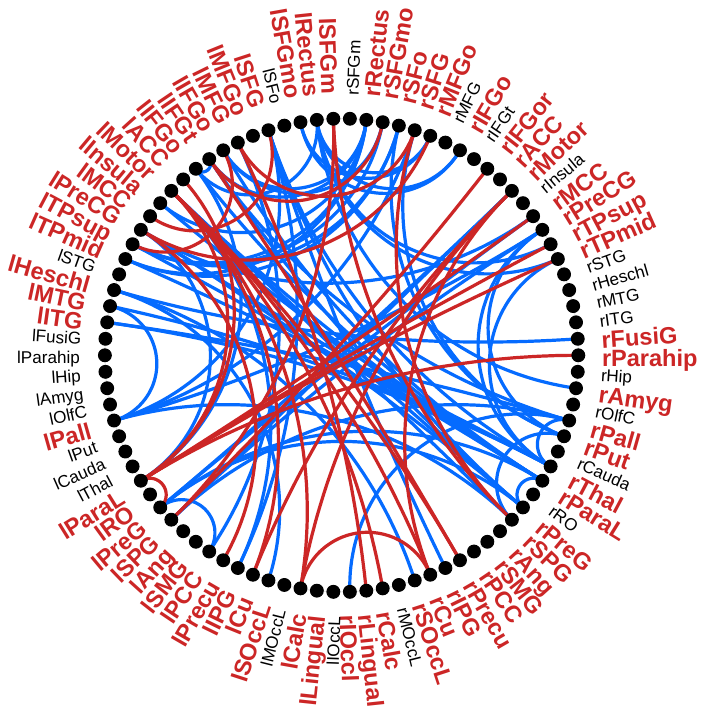

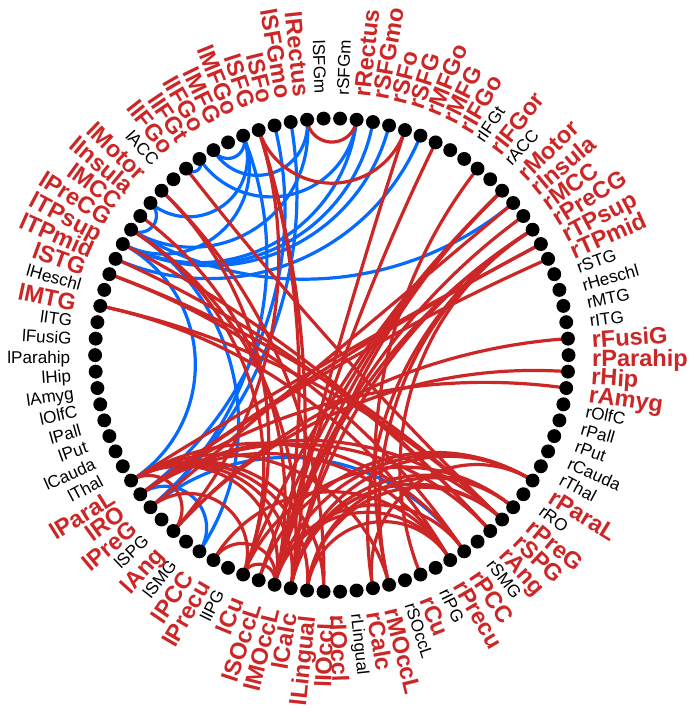

A)

**Theta Dice Score: 0.47**

B)

**Alpha Dice Score: 0.64**

C)

**High Beta Dice Score: 0.62**

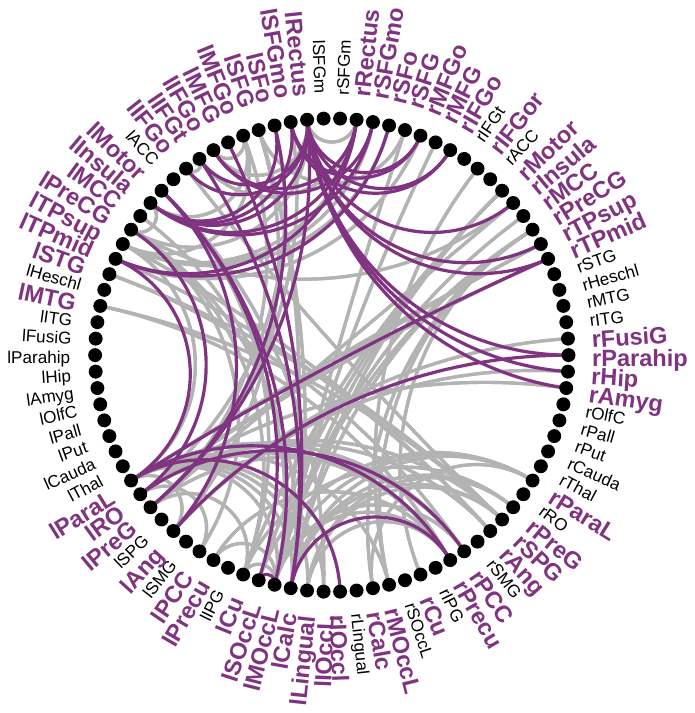

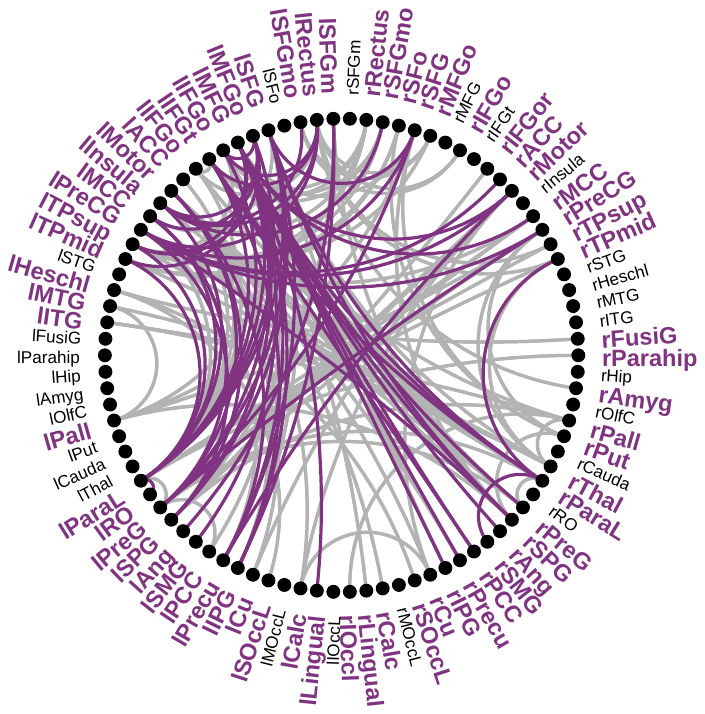

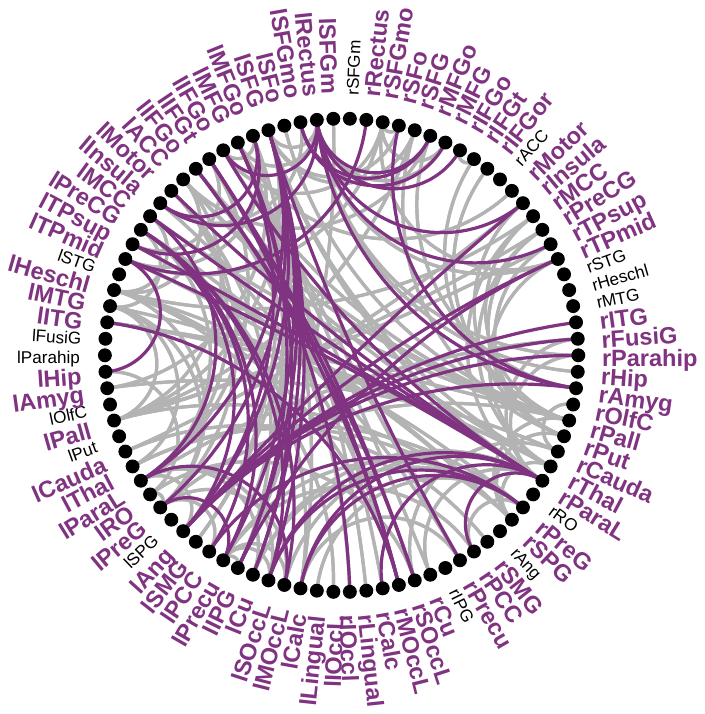

***SM 1****. Replication of FC-SAUs correlations. On the left, a summary graph of the links not shared between the networks obtained from previous study and the subsample from the present research is represented. Blue lines depict links from a previous study. Red lines depict links from the present study. On the right is a summary graph of the connections shared between both analyses. Purple lines depict shared links. Grey lines depict links from the previous study that were not replicated. A) CBPT analyses on the theta band. B) CBPT analyses on the alpha band. C) CBPT analyses on the high beta band.*

**Supplementary Material 2: Stage * Sex interactions in GLMM analyses**

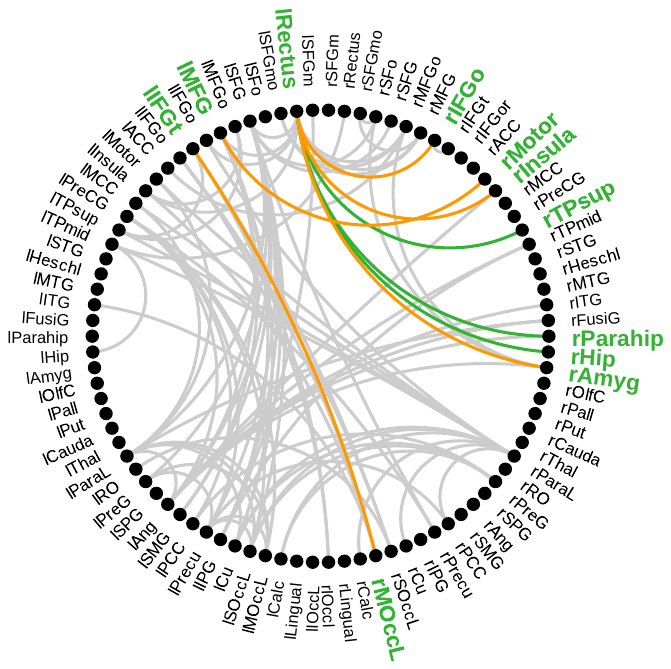

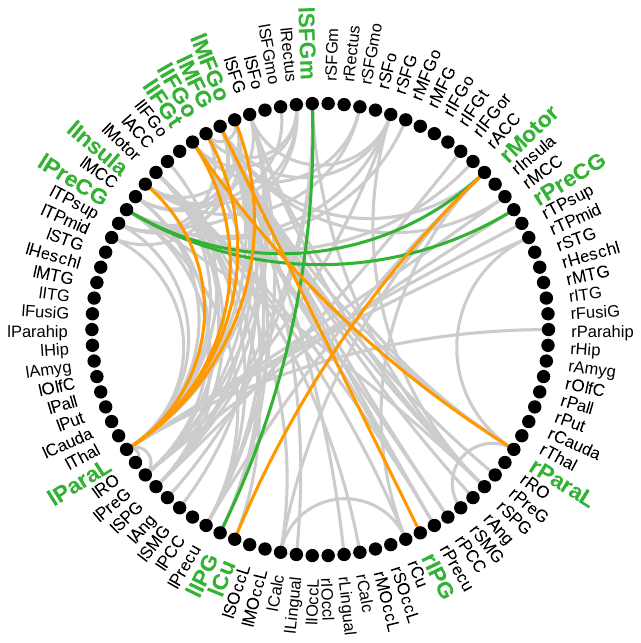

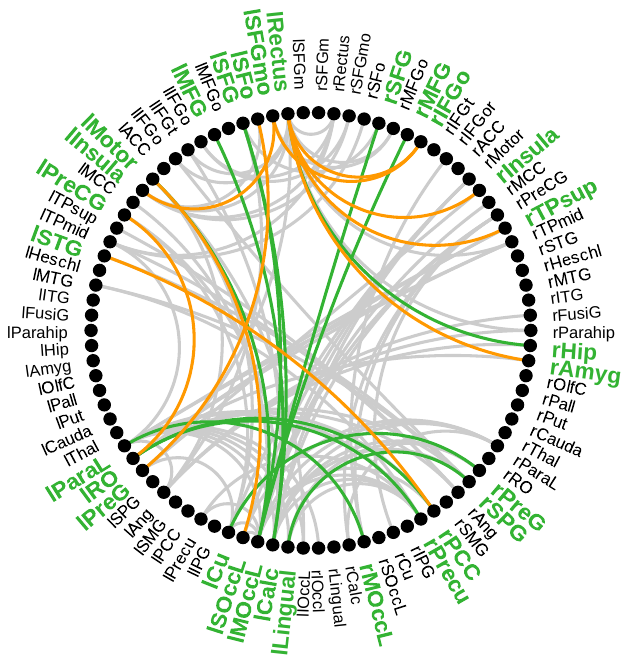

**Sex*Stage Interaction**

**C**

**A**

**B**

***SM 2****. General linear mixed model of functional connectivity in relation to biological sex throughout the study. A) Summary graph of the links with significant Sex*Stage interaction. A) Theta band network analysis. B) Alpha band network analysis. C) High beta band network analysis. Green lines represent the links with significant Sex*Stage interaction. Yellow lines represent the links with significant Sex*Stage and SAUs*Stage interaction. The grey lines represent those links of the original network that did not show significant interactions.*
